# Learning and Memory Performance in a Rat Model of Perigestational Opioid Exposure

**DOI:** 10.64898/2025.12.08.693053

**Authors:** Meghan E. Vogt, Anne Z. Murphy

## Abstract

*Rationale:* Nearly one third of women of reproductive age in the United States are prescribed opioids annually; 14% will fill an opioid prescription during pregnancy, and one in five report misuse. Opioid use during pregnancy has given rise to an increasing population of infants born with gestational opioid exposure. At school age, these children show neurodevelopmental impairment and higher rates of learning disability. Objectives: To characterize how exposure to exogenous opioids during brain development affects learning and memory performance, our lab has developed a rat model of perigestational opioid exposure that closely recapitulates a clinically relevant dosing timeline by beginning morphine exposure before pregnancy and continuing through the first postnatal week. Male and female offspring generated from this model were assessed on several facets of learning and memory during adolescence. Results: Here, we report that morphine exposure selectively impaired spatial learning, as measured by the Barnes Maze, and associative learning, as measured by an Attentional Set Shift Task, without significantly impacting working or short-term memory. Environmental enrichment rescued the spatial learning deficit in males but not females. Conclusion: Our findings suggest that gestational opioid exposure can impair performance in complex cognitive tasks that are attenuated by non-invasive, non-pharmacological intervention.

## Introduction

Over the past two decades, opioid use during pregnancy has become increasingly common in the United States. Nearly one third of reproductive-age women are prescribed opioids annually, and an estimated 14-22% fill an opioid prescription during pregnancy (Ailes et al., 2015; Ko et al., 2020). Although many of these prescriptions are intended for the management of chronic or post-surgical pain, the rise in illicit opioid use has greatly amplified the number of women using opioids during pregnancy. As opioids readily cross the placenta, gestational use impacts the developing fetus as well as the mother. Mirroring the nationwide increase in opioid use among pregnant women, the incidence of babies born with opioid withdrawal symptoms has also been increasing with the current national rate estimated at approximately 6 infants for every 1000 hospital births (Healthcare Cost and Utilizaiton Project (HCUP), 2024). The short-term withdrawal symptoms in infants are well established and typically managed postnatally via non-pharmacological (low stimulus environment) and pharmacological (oral morphine) strategies. However, a complete understanding of long-term effects of gestational opioid exposure on behavior and cognition remains elusive.

Clinically, infants with gestational opioid exposure display neurodevelopmental impairment and altered functional connectivity of the prefrontal cortex (Hunt et al., 2008; Radhakrishnan et al., 2021). Higher rates of learning disability and memory deficits have also been reported in this population at school age (Fill et al., 2018; Maguire et al., 2016; Sundelin Wahlsten & Sarman, 2013; Wilson et al., 1979). Adolescents with a history of gestational opioid exposure that performed a working memory-selective attention task during an fMRI showed working memory deficits and altered activation of the prefrontal cortex during the most challenging task versions (Sirnes et al., 2018). While clinical data is beginning to highlight the cognitive impairments of children exposed to opioids *in utero,* preclinical work characterizing the nature and severity of these deficits is far from complete.

Preclinical studies have reported that morphine administration to pregnant rats on gestational days 11 through 18 results in offspring showing mild performance deficits in the spatial water maze and radial arm maze (Ahmadalipour et al., 2018; Schrott et al., 2008; Šlamberová et al., 2001). Deficits in recognition, reference, and spatial memory were also reported in rats that were continuously exposed to either methadone or buprenorphine during gestation via osmotic minipumps (Kongstorp et al., 2020; Lum et al., 2021). More complex cognition is also reported to be negatively impacted by perigestational morphine exposure in mice, as demonstrated by reduced accuracy in the five-choice serial reaction time task. These reported deficits in executive function appear sex-specific as males display poorer performance than females and may be driven by differential microglia-dependent synaptic refinement (Smith et al., 2022; Smith et al., 2025).

Our lab has developed a perigestational opioid exposure model that aims to closely mimic an administration timeline of clinical relevance; here we use daily, intermittent morphine doses that begin before pregnancy, continue through gestation, and are tapered postnatally (Substance Abuse and Mental Health Services Administration, 2014). With this model, we have reported that perigestational morphine exposure impacts predictability of maternal care (Searles et al., 2025) and several aspects of behavior in adolescent offspring, including reduced juvenile play (Harder et al., 2023) and altered alcohol and sucrose preference (Searles et al., 2023. Changes in adult immune response have also been reported (Harder et al., 2025a, 2025b).

To further characterize the phenotype of rats with gestational opioid exposure, the present study explored performance on several learning and memory tasks in adolescence. Here, we report that morphine exposure selectively impaired spatial learning, as measured by the Barnes maze, and associative learning, as measured by an attentional set shift task, without significantly impacting working or short-term memory. Follow up studies assessed the ability of environmental enrichment to rescue the observed deficits.

## Methods

### Subjects

Female Sprague Dawley rats (Charles River Laboratories) were bred with sexually experienced males to generate offspring for experiments. Offspring were weaned at postnatal day 21 (P21) into Optirat GenII individually ventilated cages (Animal Care Systems) with corncob bedding (Bed-o’Cobs; The Andersons). Rats were housed in same-sex, same-treatment groups of two to four on a 12/12-hour light/dark cycle (lights off at 8:00 AM). Rodent chow (Laboratory Rodent Diet 5001 or 5015 for breeding pairs; Lab Diet) and water were provided *ad libitum* in the home cage. All experiments were approved by the Institutional Animal Care and Use Committee at Georgia State University and performed in compliance with the National Institutes of Health Guide for the Care and Use of Laboratory Animals. All efforts were made to reduce the number of rats used and minimize pain and suffering.

### iPrecio Pump Implantation Surgery

Female Sprague Dawley rats (P60 – P70) were implanted with programmable micro-infusion minipumps (iPrecio SMP-200; Alzet) under sterile conditions. Rats were initially anesthetized with 5% isoflurane, then maintained on 2.5-3.5% during the surgery. The right flank was shaved and prepped with povidone-iodine antiseptic scrubs and 5% povidone-iodine solution (Betadine). A 3-4cm incision was made in the skin along midline and a subcutaneous pocket was created using curved hemostats. The pre-programmed pump filled with sterile saline was implanted in the pocket and sutured to the muscle wall after which the initial incision was closed with wound clips and covered with topical bacitracin. Following surgery, each rat awoke from anesthesia in a clean recovery cage before returning to their home cage. Rats received a subcutaneous injection of 5mg carprofen per kg body weight before and twenty-four hours post-surgery. Remaining wound clips were removed two weeks post-surgery.

### Perigestational Opioid Exposure Model

Before surgical implantation, pumps were programmed to deliver 30µL of either morphine or saline across one hour three times per day (12:00 PM, 8:00 PM, 4:00 AM); during non-infusion hours the pumps were set to the minimum flowrate of 0.1µL/hour. Rats were randomly assigned to either perigestational morphine exposure (MOR) or control (VEH). One week after surgery, rats in the MOR group began receiving morphine while control dams continued to receive sterile saline. One week after morphine initiation, females were paired with sexually experienced males for two weeks to induce pregnancy. Morphine (or saline) exposure continued throughout gestation. Rats were initially administered 10mg morphine per kg body weight per day, with doses increasing weekly by 2 mg/kg/day until 16 mg/kg/day was reached (**Figure 1**). At approximately embryonic day 18, infusions decreased to twice-a-day to minimize early-life pup mortality. Post-parturition, dams continued to receive decreasing doses of morphine until P7 when administration ceased, and morphine was replaced with sterile saline. This protocol closely mirrors the clinical profile of a late adolescent female who is regularly using an opioid, becomes pregnant, and continues using throughout pregnancy and following birth.

**Figure 1:**
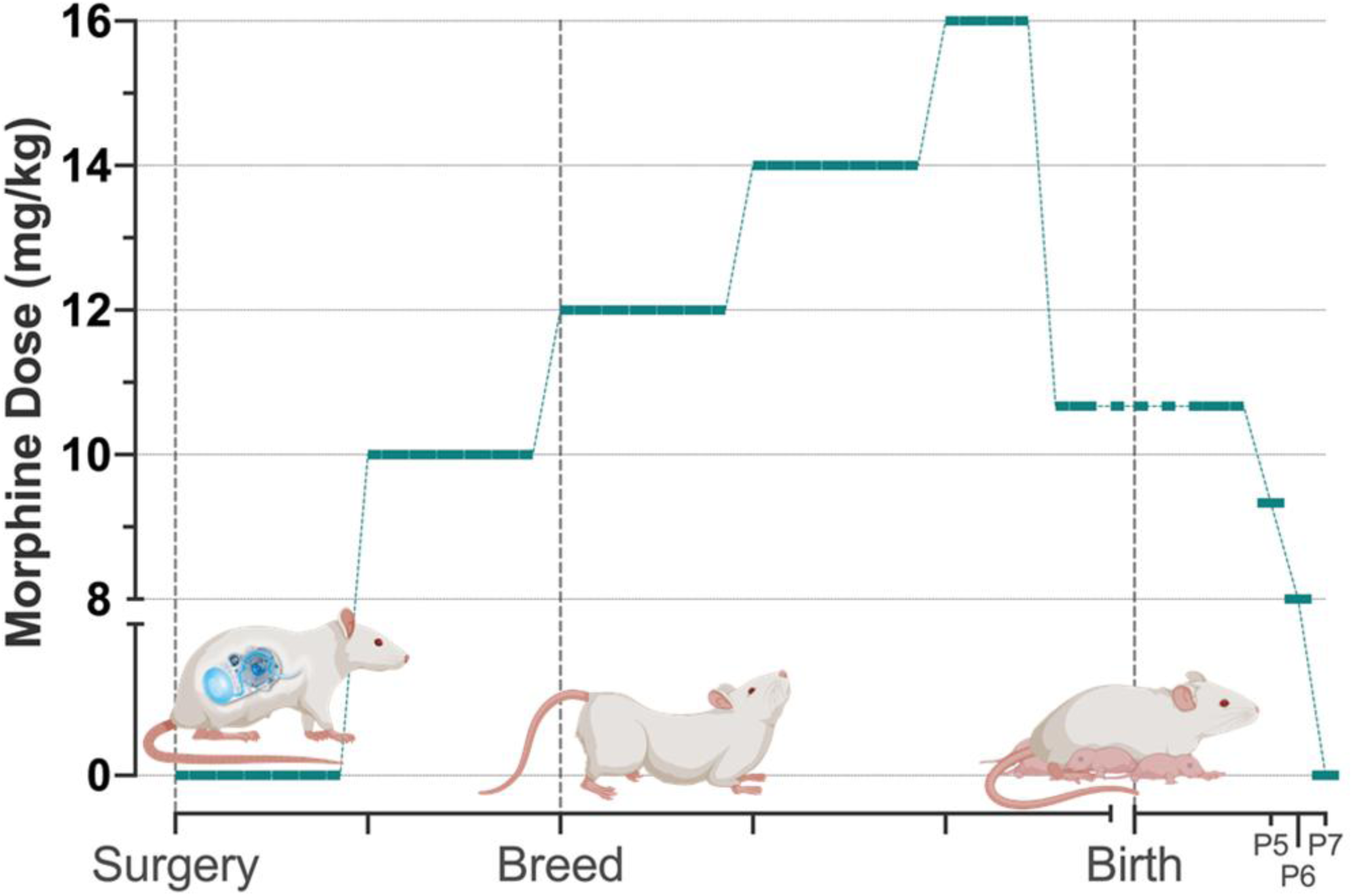
Perigestational morphine administration paradigm. Timeline of paradigm on x-axis; daily dose of morphine on y-axis. *Created in part with BioRender.com*.

### Handling & Puberty Onset

To habituate rats to the researcher and reduce stress during testing, rats were handled individually for 1-2 minutes in the home cage 3-5 times a week from weaning until onset of behavior testing. Rats were assessed for physical signs of puberty onset once per day beginning P30 for females and P40 for males. Vaginal opening was observed by applying pressure to the hindlimb area to induce dorsiflexion of the spine. In males, preputial separation was assessed by placing the rat in a supine position and gently palpating the base of the penis. The day on which full vaginal opening or full preputial separation was observed was recorded as the age of puberty onset.

### Behavioral Assessments

All testing took place during the rats’ dark cycle. Behavioral testing was initiated in adolescence to mirror clinical studies reporting impaired memory in opioid-exposed adolescents. Behavior assays varied in length, complexity, and amount of handling required. Thus, to limit stress and age confounds, all subjects underwent the tests in the same order (**Figure 2**). Due to their extensive protocols, each rat was tested on either the attentional set shift or the Barnes maze, not both. Before each task, rats were placed in the testing room (or adjacent room for Barnes Maze) for 30-60 minutes to acclimate in their home cage under dim or red lighting. All apparatuses were cleaned with 70% ethanol before and after testing. All assays were conducted around the same time of day by an experimenter blind to treatment.

**Figure 2:**
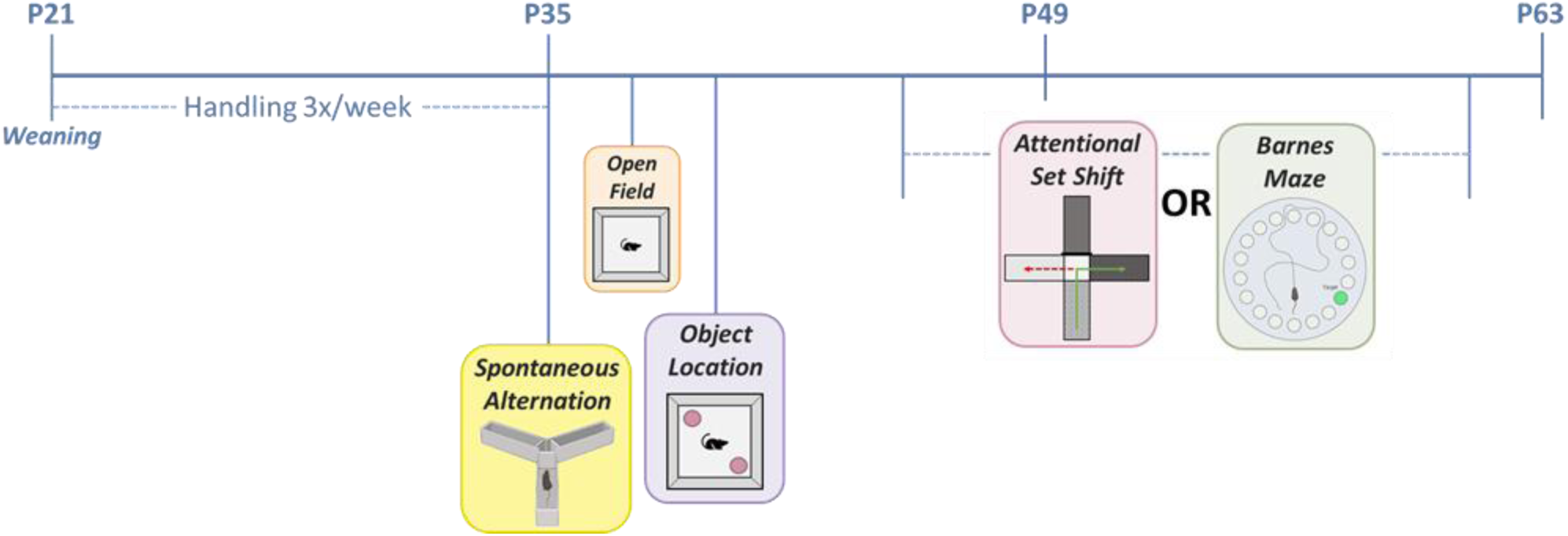
Behavior timeline.

#### Spontaneous Alternation Task

To assess spatial working memory, adolescent rats (P35-39) were tested on the spontaneous alternation task using a Y-maze (arms measuring 52cm x 12cm with 22cm walls) under dim lighting. Rats were given 5 minutes to explore the Y-maze. Behavior was videorecorded, and the following measures analyzed using ANY-maze scoring software (RRID:SCR_014289): total distance traveled, number of entries into each arm, and number of alternation paths defined as three consecutive entries into unique arms, e.g. A-B-C or C-A-B. Percent alternation was used as an indicator of working memory performance and calculated as Number of Alternation Paths / (Total Entries – 2) * 100. Rats with intact spatial memory will have percent alternation scores higher than chance.

#### Open Field

One to three days following the spontaneous alternation task (P36 – P43), rats were assessed for baseline anxiety and locomotion using the open field test. Under dim lighting, rats were given 10 minutes to explore the arena (120cm x 120cm x 40cm), and behavior was videorecorded for offline analysis using ANY-maze scoring software. The following measures were assessed: total distance traveled, mean speed, number of center entries, latency to center, and time spent in center. Rats with typical levels of anxiety will spend the majority of their time along the outer walls of the arena with minimal center time.

#### Object Location Task

To assess short-term and long-term spatial memory, rats were tested on the object location task. Rats were given a 5-minute habituation trial in an empty arena (120cm^2^) under dim lighting. For Trial 1, two identical objects – 250mL glass flasks or bottles filled with tinted water – were placed in opposite corners of the arena, 15cm from adjacent walls; rats were given 5 minutes to investigate the objects. After Trial 1, rats were returned to the home cage for a 10-minute (short-term) or 24-hour (long-term) inter-trial interval. For Trial 2, one of the objects was moved to a new corner, again 15cm from adjacent walls, and rats were given 5 minutes to investigate. Behavior during both trials was videorecorded for offline analysis. ANY-maze scoring software was used to automatically score number of investigations of each object and time spent investigating each object. The discrimination index, an indicator of short-term memory performance, was calculated using the following formula: (Time with Object in Novel Location – Familiar) / (Total Time). A positive discrimination index indicates more time spent interacting with the object in the novel location.

#### Attentional Set Shift

To assess associative learning and cognitive flexibility, adolescent rats (P45) were tested with an attentional set shift task using a modified protocol from Kougias et al. (2018). The attentional set shift was conducted in a four-arm apparatus elevated 90cm under dim lighting; each arm was defined by two dimensions – color (dark or light) and texture (smooth or rough). Prior to testing, rats were given sucrose pellets in their home cage for four consecutive days to familiarize them with the reward and encourage consumption during the task. After sucrose exposure, rats were placed in the sucrose-baited apparatus and allowed to explore for 5 minutes. Four habituation trials were run across two consecutive days. Following habituation, rats were pre-trained on the task procedure. For each pre-training trial, the rat was placed at the end of one arm (start arm) with the opposing arm blocked; the rat would leave the start arm and enter one of the adjacent arms to receive a sucrose reward. Choices were recorded to determine whether individual rats showed innate preferences for a particular arm, color, or texture. Twenty-four pre-training trials were conducted across 2 days. The apparatus was rotated 90° between each trial to discourage use of extramaze cues, and the start arm was alternated for each trial.

Set one training began 24 hours after pre-training following the same procedure but with only the correct arms resulting in a sucrose reward. The criteria assignments that defined the correct arms – light, dark, rough, or smooth – were counterbalanced, so each experimental group had representation of all criteria. If a rat showed a preference for a particular arm, color, or texture during pre-training, they were intentionally assigned a non-preferred dimension. Rats ran 20-30 trials per day until they reached performance threshold of 8 consecutive correct choices or exclusion threshold of 100 trials.

Rats that met performance threshold on set one training began set shift training 24-48 hours after reaching criterion. For set shift training, each rat had a new informative dimension assigned. For example, if the rat received a reward for entering a dark arm in set one, it would now receive a reward when it entered a rough arm. Thus, each choice was either correct (following the new rule), a perseverance error (following the set one rule), or an omission error (following neither rule). Each rat ran 60 set shift trials regardless of performance.

#### Barnes Maze

To assess spatial learning, long-term memory, and cognitive flexibility, the Barnes maze was conducted on a 120cm circular platform elevated 90cm (60270; Stoelting) under directed, bright lighting. There were twenty 10-cm holes along the edge of the platform, 19 of which were blocked by false bottoms. The 20^th^ hole led to a 20cm x 12cm x 12cm escape chamber that was baited with five 45mg sucrose pellets. Large, distinct geometric shapes cut from paper were placed on the walls surrounding the platform to act as extramaze visual cues. The Barnes maze protocol consists of four phases: acquisition, probe trial, reversal training, and reversal probe trial. Prior to testing, rats were acclimated to the adjacent room under red light, and sucrose pellets were placed in their home cages to familiarize them to the reward. Rats were also habituated to the escape chamber for 2 minutes before the first acquisition trial.

During acquisition (P45 – P49), rats ran 10 training trials across 3 days to learn the location of the escape chamber, which remained in the same location for the duration of this phase. For each trial, the rat was placed inside a 16cm x 16cm cylinder in the center of the maze for 5-10s before the cylinder was lifted, allowing the rat to explore for a maximum of 120s. If the rat entered the escape chamber within 120s, it was kept in the chamber for 30-60s before being returned to the home cage. If the rat did not enter the escape chamber within 120s, it was gently guided to the target by the researcher and kept in the chamber for 30-60s before being returned to the home cage. Rats spent 5-10 minutes in their home cage between same-day trials. Latency to reach escape chamber entrance, i.e. target, and latency to enter escape chamber were recorded live. Behavior was also videorecorded for additional offline analysis.

Seventy-two hours following the last acquisition trial (P50 – P54), rats ran the probe trial in which all 20 holes were blocked with false bottoms. As in the training trials, the rat was released from the center of the maze and given 120s to explore. Behavior was videorecorded for offline analysis.

Approximately 3 days following the probe trial (P51 – 58), the platform was rotated 180° to change the position of the escape chamber. Rats ran 6 reversal training trials across 2 days to learn the new location of the escape chamber. These training trials were run in the same manner described above for initial acquisition. Forty-eight hours following the last reversal trial (P54 – 61), rats ran another probe trial as described above.

To assess if differences in visual acuity impacted the use of the spatial cues, a separate cohort of rats (3 VEH M, 3 VEH F, 4 MOR M, 2 MOR F) was run on a landmark version of the Barnes Maze in which the location of the escape chamber varied from trial to trial and was always marked with a visual cue. Rats ran 8 total trials, and behavior was videorecorded for offline analysis.

Videos were hand-scored by a researcher blind to treatment group. The following measures were recorded: *hole deviation score –* distance (in number of holes) between the first hole visited and the target; *errors –* number of head deflections into a non-target hole before reaching the target; *perseverance errors –* number of visits to original target location during reversal training; and *strategy* used to reach target *–* serial, spatial, random, or guided.

Serial – Rat visits at least two adjacent holes consecutively before reaching target
Spatial – Hole Deviation Score of 2 or less
Random – Rat crosses center of maze between hole visits at least twice
Guided – Researcher led rat to the escape chamber (after 120s)

ANY-maze scoring software was used to score amount of time spent in each quadrant of the platform during probe trials and generate line plots of the rat’s position within the platform.

### Environmental Enrichment

Beginning P7, litters received home cage enrichment in the form of additional nesting material and shelters. In addition to home-cage enrichment, between P22 and P40, rats spent one hour per day in an enrichment arena. The enrichment arena was a 120cm x 120cm x 40cm square arena divided into two-120cm x 60cm x 40cm halves, one for each sex, that contained bedding of multiple types, nestlets, shelters, tunnels, and ping-pong balls. Eight to twelve rats of the same sex were placed into the arena together for physical, sensory, and social enrichment.

### Statistical Analyses

All data met the requirements for parametric analyses – normality and homogeneity of variance – unless otherwise noted. The effects of treatment, sex, and trial (where applicable) were assessed using t-tests (k=2), one or three-way mixed model ANOVAs (k=3) with an alpha level of 0.05 unless otherwise noted. As repeated measures ANOVA cannot handle missing values, mixed models were fit with Greenhouse-Geisser correction, when necessary, as implemented in GraphPad Prism 10.5.0. When no sex difference or sex x treatment interaction was present, sexes were combined for analysis to improve power. Tukey’s or Sidak’s post hoc tests were conducted to determine significant mean differences between *a priori* specified groups. No significant litter or cohort effects were observed when run as covariates, so offspring from 8 vehicle-exposed litters (VEH), 8 morphine-exposed litters (MOR), and 4 morphine-exposed litters that received enrichment (MOR EE), generated across 15 months, were combined into one dataset for analysis. Graphs show mean ± SEM unless otherwise noted. GraphPad Prism 10.5.0 was used for all statistical analyses.

## Results

### Puberty Onset

To determine if morphine exposure influences puberty onset, we investigated timing of vaginal opening in females and preputial separation in males. Vaginal opening in females occurred approximately one week prior to preputial separation in males (P35 vs P42, respectively) [F_sex_(1,39)=323.3, p<0.0001]. Perigestational morphine exposure significantly delayed puberty onset in females by two days (P34 vs P36), and no significant impact of treatment was observed in males [F_treatment x sex_F(1,39)=7.652, p=0.0086; **Figure 3**].

**Figure 3:**
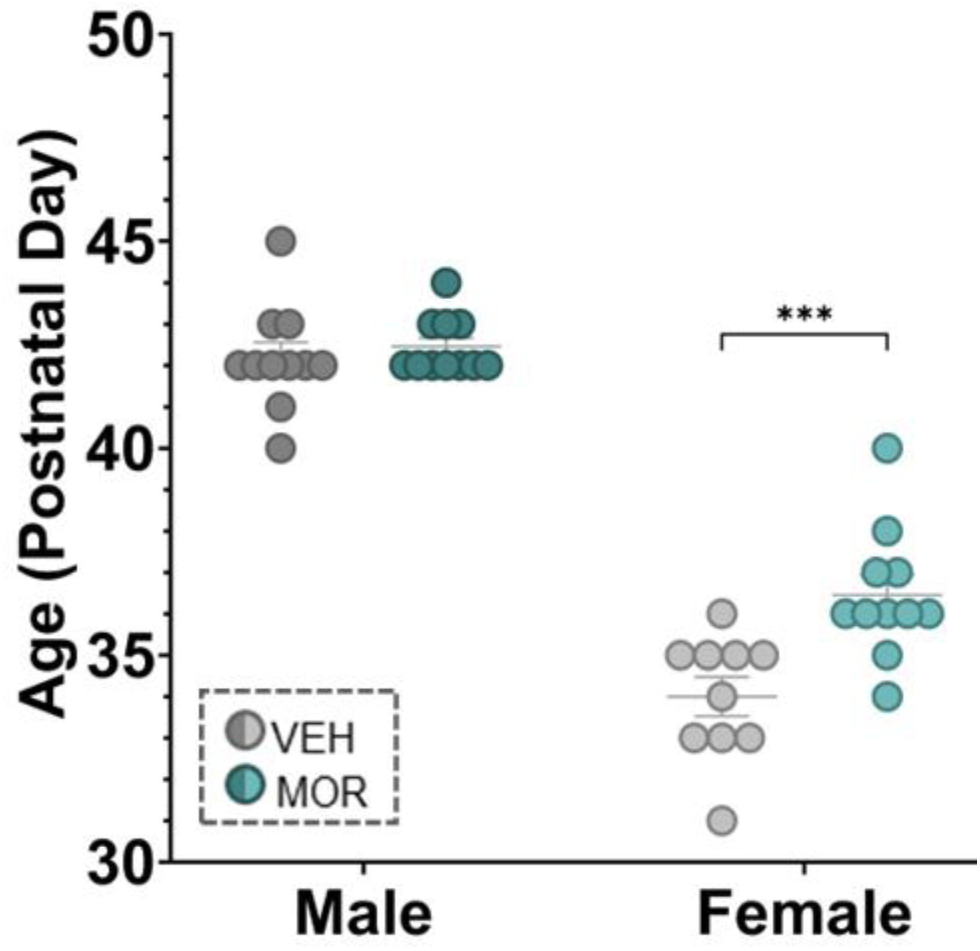
Mean age of puberty onset [n=10-11 per sex per treatment]. Males are darker shades; females are lighter. ***p<0.0001.

### Behavioral Assessments

#### Spontaneous Alternation Task

To investigate working memory, adolescent rats were tested on the spontaneous alternation task. When placed in the 3-armed apparatus, rats with intact working memory will spontaneously alternate the arms they visit such that they enter each arm once before re-entering another. Working memory performance was quantified as the percentage of alternation paths made (consecutive entries into 3 unique arms) from the total number of possible alternation paths, as determined by total arm entries. Three subjects (1 VEH F, 1 MOR M, 1 MOR EE F) were excluded from analysis for making less than 10 total arm entries during the 5-minute task. No effects of morphine exposure were observed on percent alternation, with VEH and MOR showing scores of 73.77% and 75.36%, respectively. Environmental enrichment did not significantly impact percent alternation, with MOR EE rats averaging 71.25%. [F_treatment_(2,57)=0.7597, p=0.4725; **Figure 4a**]. However, MOR EE rats displayed an increased tendency to escape the apparatus during testing, as reflected in a significant treatment effect on total number of arm entries [F_treatment_ (2,57)=5.058, p=0.0095; MOR EE vs VEH, p=0.0204; vs MOR, p=0.0201; **Figure 4b**]. Notably, all rats that escaped during testing were immediately placed back into the arm from which they escaped and allowed to complete the task.

**Figure 4:**
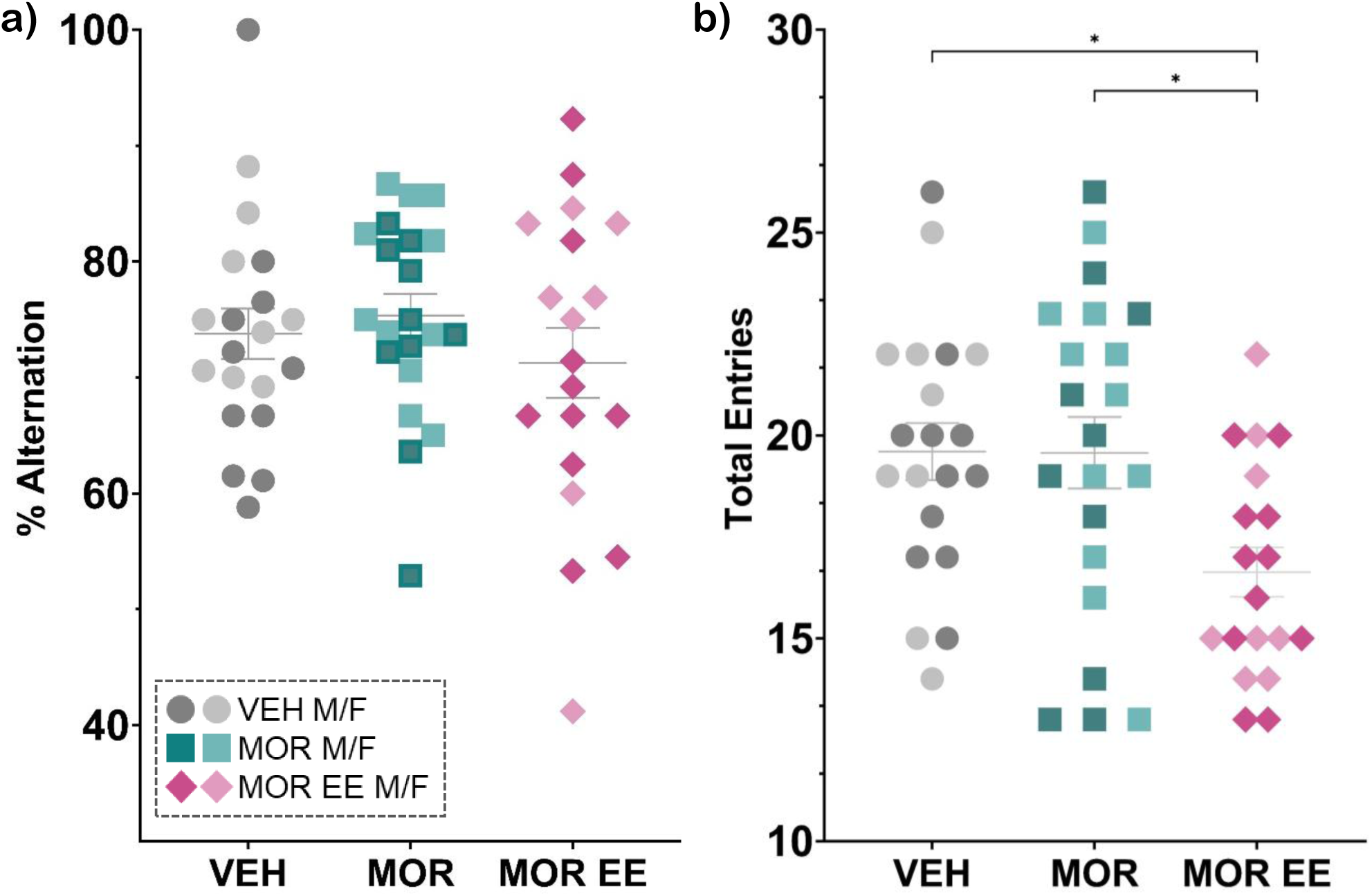
Percent alternation (a) and total entries (b) during spontaneous alternation task [n=8-11 per sex per treatment]. Males are darker shades; females are lighter. *p<0.05

#### Open Field

At P36 – P43, rats were assessed for possible treatment effects on baseline anxiety and locomotion. Our analysis showed no effect of morphine or enrichment exposure on time spent in center [F_treatment_(2,61)=0.2040, p=0.8160; **Figure 5a**] nor on distance traveled [F_treatment_ (2,61)=0.6657, p=0.5176; **Figure 5b**] during the 10-minute observation.

**Figure 5:**
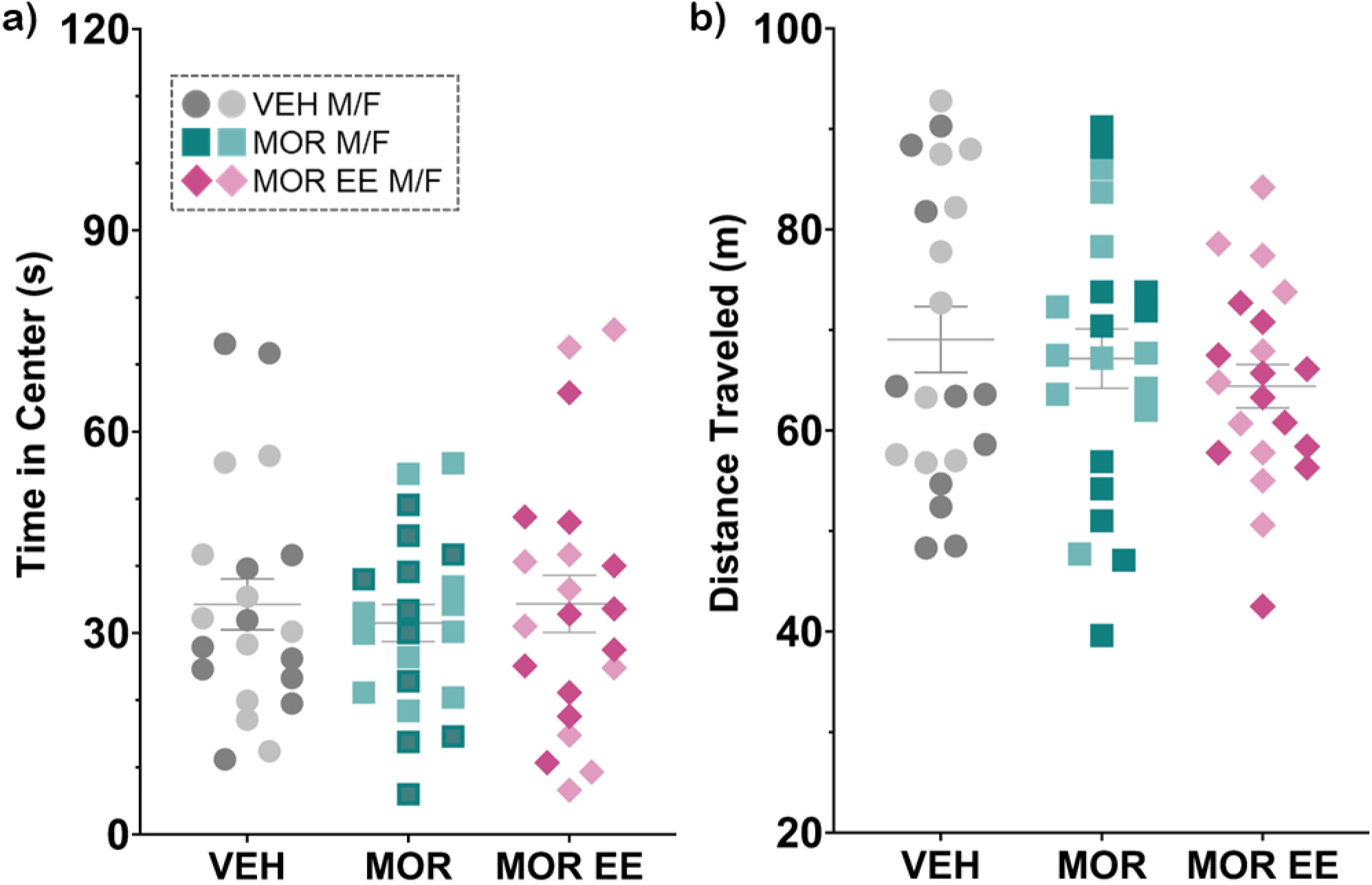
Open field anxiety-like behavior (a); locomotion (b) [n=10-11 per sex per treatment]; Males are darker shades; females are lighter.

#### Object Location Task

To investigate short-term and long-term spatial memory, the object location task was conducted with a 10-minute and 24-hour inter-trial interval (ITI), respectively. Total time spent investigating objects was reported as a control measure. To quantify memory performance, a discrimination index (DI) was calculated for each rat for trial 2; a higher DI signifies larger proportion of time spent with object in novel location, i.e. better memory of object in familiar location. For the short-term object location task (10-minute ITI), no significant treatment effects were found in total investigation time [F_treatment_(1,40)=0.4328, p=0.5144; **Figure 6a**]. However, both VEH and MOR groups spent significantly less time investigating objects during trial 2 (familiar or novel location) [F_trial_(1,40)=22.73, p<0.0001; VEH T1 vs T2, p=0.0002; MOR T1 vs T2, p=0.0133]. VEH and MOR discrimination indexes did not significantly differ from each other [t(40)=0.6215, p=0.5378; **Figure 6b**] nor from zero [VEH t(19)=1.720, p=0.1016; MOR t(21)=1.168, p=0.2559].

**Figure 6:**
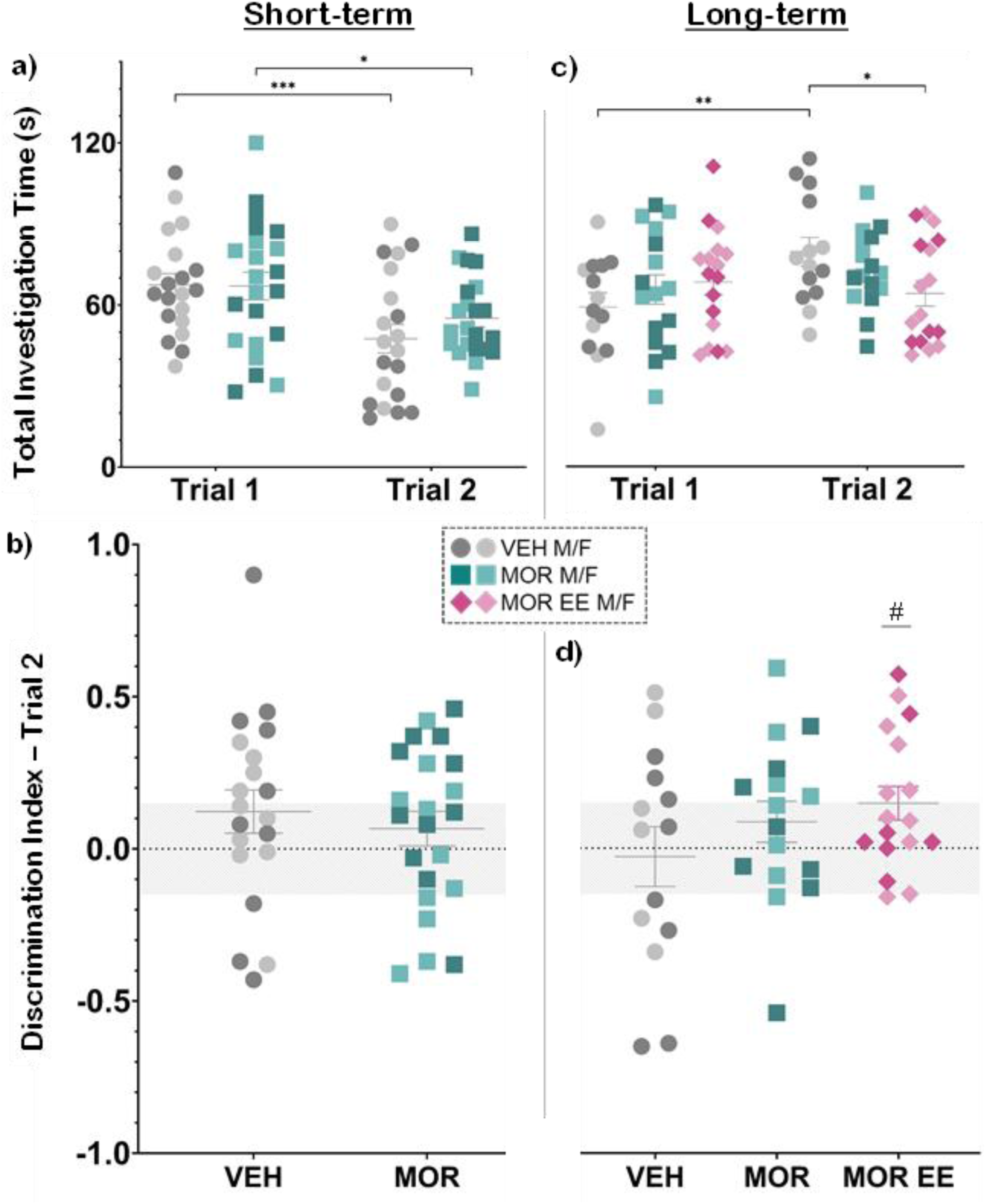
Total time spent investigating objects and trial 2 discrimination index for short-term (a, b) and long-term (c, d) object location task [short-term n=10-11 per sex per treatment; long-term n=6-10 per sex per treatment]. Males are darker shades; females are lighter. *p<0.05; **p<0.01; ***p<0.0001; # significantly (p<0.05) different from 0.0

A separate cohort of adolescent rats was assessed for long-term memory performance using the 24-hour ITI. Here, we observed a significant main effect of trial [F_trial_(1,44)=5.726, p=0.0210], with trial 2 total investigation time of both objects slightly higher than trial 1, and a significant trial x treatment interaction in total investigation time [F_trial x treatment_(2,44)=4.695, p=0.0142]. Post-hoc analyses revealed that VEH rats spent significantly more time investigating either object during trial 2 [T1 vs T2, p=0.0013]. However, even with an overall increase in total investigation time during trial 2, neither VEH nor MOR rats had discrimination indexes that were significantly different from zero [VEH t(13)=0.2842, p=0.7807; MOR t(15)=1.280, p=0.2199].

We next assessed whether environmental enrichment impacted long-term spatial memory and found that MOR EE rats, similar to MOR, spent less time investigating objects compared to VEH during trial 2 [T2 VEH vs MOR EE, p=0.0288; **Figure 6c**]. The discrimination index of MOR EE rats significantly differed from zero, however, they also had low scores with a mean of only 0.1471 [MOR EE t(16)=2.667, p=0.0169; **Figure 6d**]. Together, these data suggest that the task may have been too challenging for the rats to successfully perform at this age of early adolescence or, alternatively, that the objects used did not elicit enough engagement to be memorable.

#### Attentional Set Shift

We next investigated the impact of perigestational morphine exposure on associative learning, which is the ability to pair two stimuli, using an attentional set shift task. In this task, rats are faced with two-arm choices and trained to select arms of a specific color or texture (stimulus 1) to earn a sucrose reward (stimulus 2) (**Figure 7a**). Rats were trained until they reached performance threshold of 8 consecutive correct choices or exclusion threshold of 100 trials. Our analysis showed no significant treatment effect in the percentage of correct choices made [t(25)=0.8448, p=0.4062]. However, only VEH rats performed significantly better than chance [VEH t(12)=2.350, p=0.0367; MOR t(13)=2.131, p=0.0527; **Figure 7b**]. MOR rats also ran significantly more trials before reaching criteria compared to VEH rats [Ⴟ_VEH_=77.8, Ⴟ_MOR_=89; U=53.5, p=0.0340; Data not shown].

**Figure 7:**
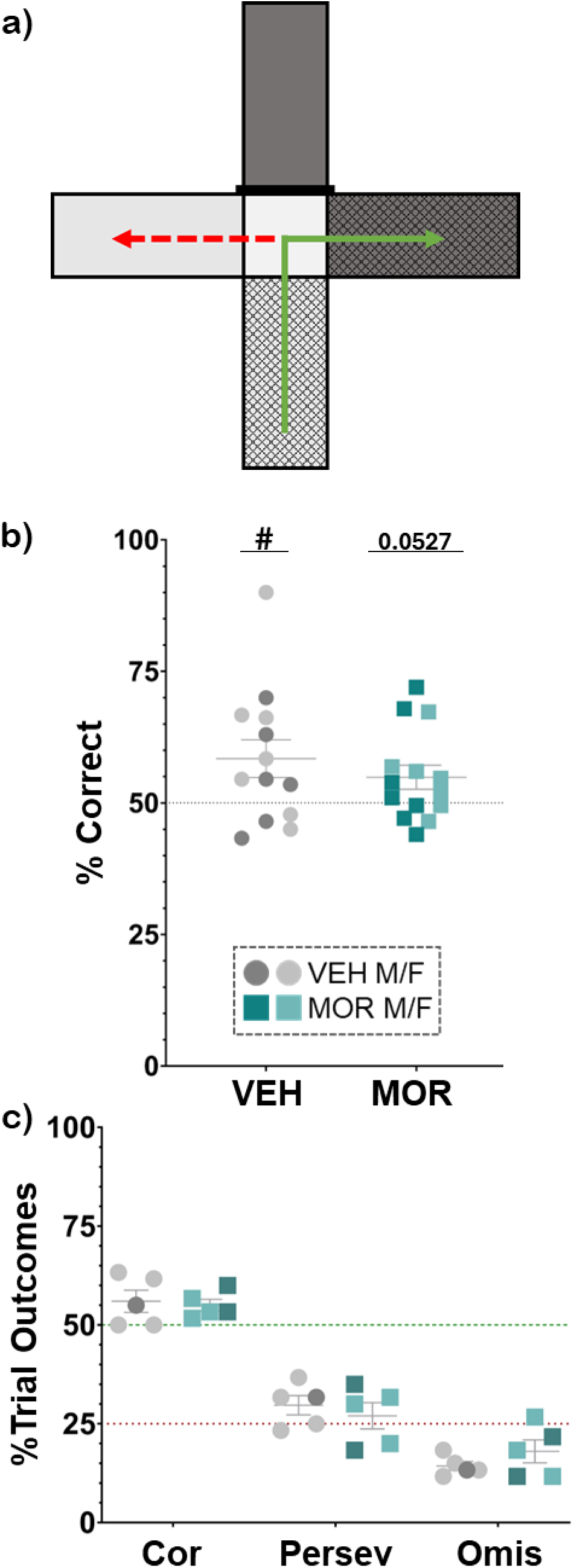
Attentional set shift apparatus with correct choice example of rough arm denoted by green arrow and omission error of light arm denoted by red dotted arrow (a). Percentage of correct choices during set one training (b). Breakdown of trial outcomes during set shift training after informative dimension (color or texture) was changed (c) [n=6-7 per sex per treatment]. Males are darker shades; females are lighter. *p<0.05; # significantly (p<0.05) different from 50%

As the attentional set shift task is quite challenging for adult rats, let alone adolescents, only 5 rats from each treatment group reached performance threshold and graduated to set shift training for assessment of cognitive flexibility; the remaining 8 VEH and 9 MOR rats were removed from further testing. Overall, MOR rats required, on average, 66 trials to reach criterion compared to 43 trials required by the VEH group, a 53% increase [data not shown]. Twenty-four hours after reaching performance threshold, set shift training began in which each rat ran 60 trials with a new informative dimension, *i.e.* if an arm color (*e.g.* dark) informed rewards originally, now a texture (*e.g.* rough) was correct. Thus, each choice the rat made was now either correct, previously correct (perseverant), or never correct (omission). No treatment effects were observed in distribution of choices [% Correct: 56 VEH vs 55 MOR; % Perseverance: 30 VEH vs 27 MOR; % Omission: 14 VEH vs 18 MOR; **Figure 7c**].

#### Barnes Maze

For the final task, rats were assessed for spatial learning using the Barnes maze. Here, rats were given 10 trials to learn the location of an escape chamber among 19 false exits on an elevated platform illuminated by mildly aversive bright light (see **Figure 8a**). All rats successfully learned the escape location as demonstrated by a significant decrease in latency to reach the target across trials with no effect of treatment [F_treatment_(2,48)=0.9380, p=0.3985; F_trial_(4.467,211.9)=21.06, p<0.0001; **Figure 8b**]. In early trials, all rats tend to use a serial search strategy to find the target location in which they check several, consecutive false exits before reaching the escape chamber, depicted by the line plot in **Figure 8c**. As rats become more familiar with the task, they tend to adopt the more efficient spatial strategy in which extra-maze cues are used to remember the relative location of the target, thus bypassing the non-exit holes as depicted by the line plot in **Figure 8d**. Overall, MOR rats were found to use the spatial strategy significantly less than VEH rats – an effect that was partially rescued by enrichment in males but not females [Males: F_treatment_(2,18)=5.675, p=0.0123; VEH vs MOR, p=0.0586; VEH vs MOR EE, p=0.7391; MOR EE vs MOR, p=0.0126; Females: F_treatment_(2,18)=20.65, p<0.0001; VEH vs MOR, p=0.0007; VEH vs MOR EE, p<0.0001; MOR EE vs MOR, p=0.2601 **Figure 8e**]. To ensure decreased use of the spatial was not due to visual impairment, we ran a landmark version of the task in which the target location changed with each trial and was always marked with a visual landmark cue and found no treatment differences; VEH and MOR rats all decreased their latency to target over time and at least 65% of each group used the visual strategy by trial 7 [data not shown].

**Figure 8:**
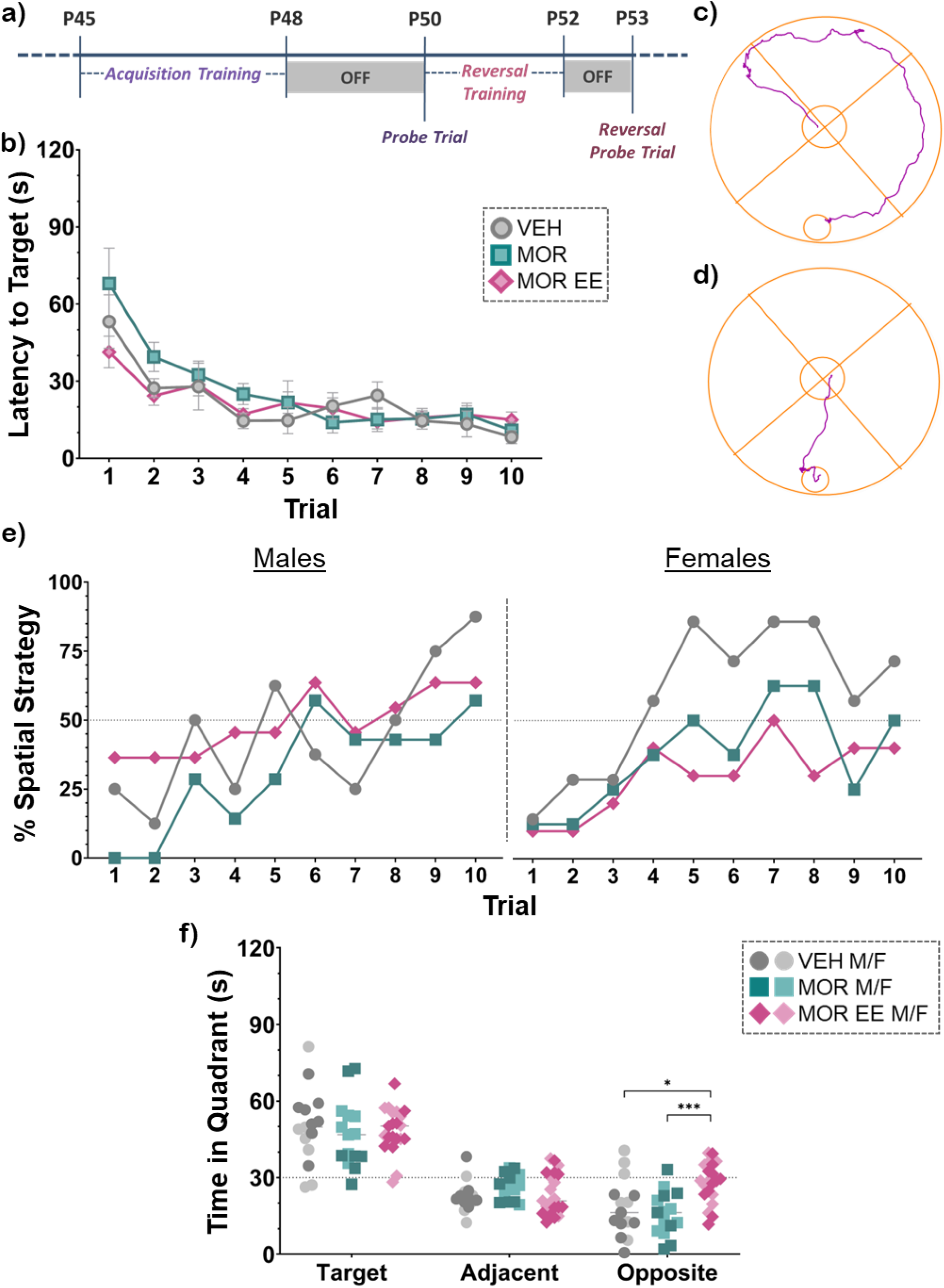
Timeline of the full Barnes Maze protocol with approximate ages (a). Latency to reach target during 10 trials of acquisition training with males and females combined (b). Examples of paths taken by rats using a serial strategy (c) vs a spatial strategy (d). Percentage of rats in each group for each trial that used a spatial strategy (e). Time spent in each quadrant type during the probe trial (f); adjacent value is mean time spent in both adjacent quadrants. [n=7-11 per sex per treatment] *p<0.05; ***p<0.0001

Seventy-two hours following acquisition, rats were tested on a 2-minute probe trial in which the escape chamber was replaced with a false bottom to assess their long-term spatial memory. Rats that remembered the location of the target should spend more time in the quadrant where it was located. Indeed, all three groups spent significantly more time in the target quadrant than either adjacent quadrants or opposite quadrant [F_quadrant_(1.563,75.04)=88.91, p<0.0001; **Figure 8f**]. Unexpectedly, MOR EE rats spent significantly more time in the opposite quadrant compared to both VEH and MOR [Ⴟ_VEH_=18.27s, Ⴟ_MOR_=15.79s, Ⴟ_MOREE_=28.09s; MOR EE vs VEH, p=0.0211; vs MOR, p=0.0004].

Approximately 24 hours following the probe trial, rats were assessed for cognitive flexibility by changing the location of the escape chamber relative to the extra-maze cues. Rats were given 6 trials to learn the new location. Visits to the original target location were scored as perseverance errors. Overall, MOR rats had a higher number of perseverance errors across the entire training (Ⴟ_VEH_=1.6, Ⴟ_MOR_=2.533), although this was not statistically significant [F_treatment_(2,48)=2.367, p=0.1046; VEH vs MOR, p=0.0858; **Figure 9a**]. All groups successfully learned the new location, as shown by a significant decrease in latency to target across trials [F_trial_(2.346,112.6)=36.63, p<0.0001; **Figure 9b**].

**Figure 9:**
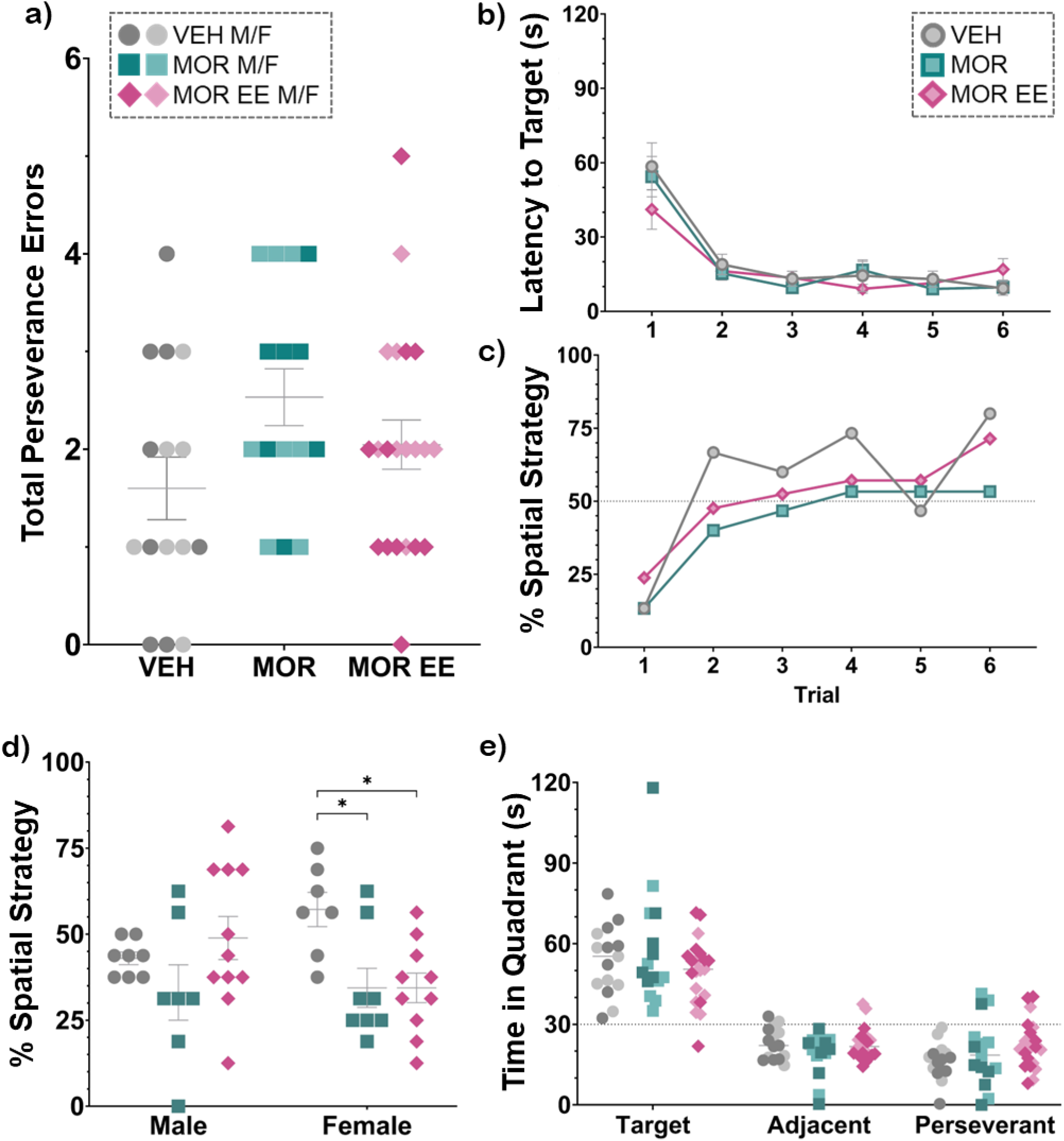
Total number of perseverance errors made across 6 reversal trials (a). Latency to reach target during 6 trials of reversal training with males and females combined (b). Percentage of rats in each group for each trial that used a spatial strategy (c). Percentage of trials that an individual subject used a spatial strategy across all 16 training trials (d). Time spent in each quadrant type during the reversal probe trial (e); adjacent value is mean time spent in both adjacent quadrants. [n=7-11 per sex per treatment] *p<0.05

Similar to the acquisition phase, MOR rats employed the spatial strategy significantly less during reversal training; this effect was partially rescued by EE [F_treatment_ (2,10)=4.196, p=0.0475; VEH vs MOR, p=0.0405; VEH vs MOR EE, p=0.5374; MOR vs MOR EE, p=0.2272; **Figure 9c**]. We further analyzed strategy usage of each individual rat by combining performance across 16 trials (10 acquisition plus 6 reversal) and analyzing the percentage of trials each subject used a spatial strategy. This analysis allows for visualization of the individual variability hidden in the collapsed group format. Our analysis showed a significant main effect of treatment and a treatment by sex interaction [F_treatment_(2,45)=3.837, p=0.0289; F_treatment x sex_(2,45)=3.523, p=0.0379; **Figure 9d**]. Post-hoc comparisons within the sexes show a significantly higher use of the spatial strategy in VEH females compared to both MOR and MOR EE females [Females: VEH vs MOR, p=0.0241; VEH vs MOR EE, p=0.0170]. Comparisons in males suggest the spatial deficit in the male MOR rats was partially rescued by EE as VEH males averaged 43% spatial usage compared to 33% in MOR males and 49% in MOR EE males, although these comparison do not reach statistical significance.

Forty-eight hours following the last reversal training trial, rats were again given a 2-minute probe assessment with no escape chamber present. Mirroring results from the original probe trial, all groups similarly spent significantly more time in the target quadrant, indicating no deficits in long-term spatial memory [F_treatment_(2,48)=0.7913, p=0.4591; F_quadrant_(1.491,71.57)=101.3, p<0.0001; **Figure 9e**].

## Discussion

The current study sought to characterize the consequences of perigestational morphine exposure on adolescent male and female rats across a battery of behavioral tests assessing multiple facets of learning and memory. We also examined the potential of environmental enrichment (EE) to rescue observed deficits. Here, we report that perigestational morphine exposure lead to delayed puberty in females and impaired associative and spatial learning in both males and females. We further show that memory and cognitive flexibility in adolescence were relatively unaffected by our morphine exposure protocol. Lastly, we show that environmental enrichment was able to rescue the spatial learning deficits in males but not females.

In the present study, we found that perigestational morphine delayed puberty in females by two days, which contrasts with a previous study that reported mild acceleration of vaginal opening in female rats administered morphine twice daily from gestational day (G) 11 through G18 (Vathy et al., 1985). Other forms of early-life adversity, such as limited bedding and exposure to environmental toxins, have also been shown to delay vaginal opening in rodents (Manzano Nieves et al., 2019; Yang et al., 2015). Reproductive behaviors, including vaginal opening, are mediated by GnRH and kisspeptin neurons of the hypothalamic-pituitary-gonadal axis; these cells express opioid receptors and, thus, can be modulated by endogenous and exogenous opioid exposure (Campideli-Santana et al., 2025). Notably, prenatal morphine exposure has been shown to alter μ-opioid receptor density in the hypothalamus, though not specifically on GnRH or kisspeptin neurons (Šlamberová et al., 2005). Together, these studies underscore the sensitivity of the developing HPG axis to opioid exposure and highlight the need to delineate the mechanisms whereby perigestational opioid exposure disrupts timing of puberty onset.

To provide a comprehensive analysis of the impact of perigestational morphine on cognitive function, we employed a battery of behavioral tests to assess distinct facets of learning and memory, including working, short-term, and long-term spatial memory; spatial and associative learning; and cognitive flexibility. We found no treatment effects on the spontaneous alternation task, suggesting no deficits in working memory. Indeed, although there was high individual variability across treatments, all subjects were able to do the task well with group means above 70% alternation and few subjects performing below 60%. As this simple task exploits the natural curiosity of rats and requires no training, a lack of treatment effect is not surprising.

In contrast to their working memory performance, both vehicle- and morphine-exposed rats performed poorly on the object location task, and no treatment effect was observed. Importantly, given the high variability of our data, the conclusion of no effect on short-term and long-term memory measured by this task is limited in strength and requires further refining and replication. A recent study by Sahid et al. (2025) reported deficits in the object location task at P28 in rats following perinatal exposure to methadone or buprenorphine. Adult rats prenatally exposed to methadone or buprenorphine also show impaired performance in the similar, yet notably nonspatial, novel object recognition task (Kongstorp et al., 2020).

As no deficits in spatial memory were observed in the object location task, we next assessed non-spatial associative learning and cognitive flexibility with an attentional set shift task. We report impairment in associative learning following our perigestational opioid exposure paradigm, with morphine-exposed rats requiring significantly more trials to reach performance threshold when trained to associate maze arms with sucrose. Our findings align with those of Kongstorp et al. (2020) in which nonspatial learning deficits were observed following prenatal exposure to methadone or buprenorphine, demonstrated by increased number of trials required to consistently pair one of three cylinders to a water reward.

Recently, our lab found that perigestational morphine exposure reduces sucrose preference in offspring (Searles et al., 2025). This observed anhedonia suggests that morphine-exposed rats may have diminished reward motivation, potentially due to altered mesolimbic dopaminergic or opioidergic signaling, that could have impacted their performance in the associative learning task. However, because sucrose was also used as a motivator for the Barnes maze, where all groups performed similarly well, it is unlikely that decreased sucrose responsiveness alone accounts for the observed associative learning deficits. Notably, the Barnes maze also utilizes avoidance-based motivation through exposure to mildly aversive stimuli (open space and bright light), whereas sucrose served as the sole reinforcer for the associative learning task. Future studies should employ associative learning paradigms that integrate both appetitive and aversive components to better understand how perigestational opioid exposure alters motivation and learning through dysregulation of reward and stress circuitry.

As no deficits in spatial working memory were observed in the simple spontaneous alternation task, we next examined whether more complex spatial and learning are affected by perigestational opioid exposure using the Barnes Maze. Performance on probe trials conducted 48-72 hours post-training revealed no effect of treatment on spatial memory as both VEH and MOR rats successfully learned and recalled the target location. However, analysis of search strategies revealed that morphine-exposed rats relied less on spatial strategies and more on inefficient serial searching, an effect that was particularly evident in females. Notably, environmental enrichment, consisting of enhanced sensory stimulation in the home cage and seven one-hour sessions in an enrichment arena during early adolescence, restored spatial strategy use in males but not females.

Our behavioral findings are consistent with prior studies reporting that prenatal opioid exposure disrupts hippocampal development and associated navigation strategies (Kongstorp et al., 2020; Šlamberová et al., 2001). Similar to the present results, Ahmadalipour et al. (2018) reported that rats exposed to morphine prenatally (G11-G18) via twice daily injections exhibited spatial learning deficits in the spatial water maze during late adolescence (P51-55). Interestingly, in their study, postnatal exercise and environmental enrichment restored spatial performance in both males and females, in contrast to our finding that enrichment rescued spatial strategy use in males only. This discrepancy may reflect the greater intensity and duration of their enrichment paradigm, which included daily treadmill exercise and continuous exposure to enriched housing from P21 through P50. Together, these studies suggest that the female brain may be less responsive or require more prolonged and/or complex forms of stimulation to achieve full recovery of hippocampus-dependent cognitive function following perigestational opioid exposure.

Current recommendations for treating infants with *in utero* opioid exposure encourage non-pharmacological interventions that strengthen the mother-infant dyad and support continued treatment following hospital discharge (Anbalagan et al., 2025; Goyal et al., 2020; Wiles et al., 2014). In line with these translational goals, incorporating environmental enrichment into preclinical models is a top priority. As noted above, Ahmadalipour et al. (2018) reported that combined home cage enrichment and daily exercise successfully reversed behavioral deficits in rats exposed prenatally to morphine. Similarly, Alipio et al. (2022) reported hyperactivity and sensory maladaptation in adolescent mice (P45 – P47) prenatally exposed to fentanyl, both of which were ameliorated by home cage enrichment from P21 and P45. Together, these studies underscore the therapeutic potential of non-invasive, non-pharmacological enrichment interventions to improve behavioral outcomes following developmental opioid exposure. Such approaches can inform clinical strategies that incorporate sensory stimulation and enrichment in the neonatal intensive care unit and home environment to promote optimal neurodevelopment in opioid-exposed infants.

Clinically, children with a history of prenatal opioid exposure have higher rates of learning disabilities and are at greater risk of poor academic performance compared to unexposed peers (Fill et al., 2018; Maguire et al., 2016; Oei et al., 2017). However, additional risk factors, such as maternal polysubstance use, confound the direct role of opioid exposure in observed negative outcomes (Tobon et al., 2019). Indeed, 39-56% of women who use opioids during pregnancy also report using tobacco and 32-50% report drinking alcohol (Jarlenski et al., 2017; Nguyen et al., 2023). Use of either of these substances alone is known to cause serious health outcomes for the fetus and neurodevelopment impairments and cognitive deficits later in life (Mattson et al., 2019; Wells & Lotfipour, 2023). Heavy smoking, in particular, also results in higher severity of Neonatal Abstinence Syndrome in newborns, requiring longer hospital stays and more extensive treatment (Jones et al., 2013). Furthermore, children exposed to opioids *in utero* have higher rates of attention disorders that likely contribute to their observed learning deficits (Nygaard et al., 2016). These statistics highlight the complex interplay between prenatal opioid exposure and comorbid maternal risk factors, underscoring the need for preclinical models that can isolate the specific neurodevelopmental consequences of opioids from those of other substances.

In conclusion, our findings indicate that perigestational morphine exposure produces specific impairments in spatial and associative learning in adolescent male and female rats while leaving general spatial memory intact. Environmental enrichment partially mitigated these effects, particularly for spatial strategy use in males. These results underscore the long-term neurodevelopmental risks of opioid use during pregnancy and highlight environmental enrichment as a promising non-pharmacological approach to support cognitive recovery in offspring exposed to opioids *in utero*.

The authors declare no competing financial interests.

## Acknowledgements

This work was supported by the National Institutes of Health (AZM; 1R21DA058889) and Georgia State University’s Brains and Behavior Program (MEV). Funding sources were not involved in study design, data collection, analysis or interpretation, manuscript writing, or the decision to publish this article. We thank the National Institute on Drug Abuse Drug Supply Program for providing morphine sulfate. We also thank AB Corley for construction of behavioral apparatuses.

## Notes

### Competing Interest Statement

The authors have declared no competing interest.

